# Enabling Organ- and Injury-Specific Nanocarrier Targeting via Surface-Functionalized PEG-b-PPS Micelles for Acute Kidney Injury

**DOI:** 10.1101/2025.07.15.664769

**Authors:** Boaz Y. Bishop, Swagat H. Sharma, Ratnakar Tiwari, Simseok A. Yuk, Sultan Almunif, Susan E. Quaggin, Evan A. Scott, Pinelopi P. Kapitsinou

## Abstract

While nanomedicine holds great promise for kidney disease, targeted delivery remains a major challenge. Most nanocarriers rely on passive accumulation or epithelial-specific ligands, limiting their utility in complex, inflamed renal environments. In acute kidney injury (AKI), inflammation and vascular dysfunction play central roles, yet targeting strategies beyond the tubule remain underexplored. Here, dual-ligand micelles are developed to enhance nanocarrier localization to the inflamed kidney by simultaneously engaging both organ- and injury-specific cues. Poly(ethylene glycol)-block-poly(propylene sulfide) (PEG-b-PPS) micelles were engineered to display two peptide ligands: CLPVASC, which preferentially distributes to the kidney, and CYNTTTHRC, which binds selectively to inflamed endothelium. These targeting motifs were incorporated via lipid-anchored peptide amphiphiles, enabling modular surface functionalization without disrupting micelle morphology, size, or charge. In vitro, dual-targeted micelles demonstrated enhanced uptake by human endothelial cells exposed to hypoxia-reoxygenation. In vivo, following unilateral renal ischemia-reperfusion injury (IRI) in mice, targeted micelles achieved selective accumulation in the injured kidney, outperforming both non-targeted controls and contralateral kidneys. Off-target distribution to liver, lung, and spleen was markedly reduced, confirming the spatial precision of the dual-ligand approach. This strategy offers a scalable, modular, and biologically informed platform for precision delivery in AKI and related inflammatory conditions.

## Introduction

Acute kidney injury (AKI), characterized by rapid decline in kidney function, occurs in approximately 7 to 18% of all hospitalized patients and up to 67% of critically ill patients ^1–3^. AKI is strongly linked to increased mortality, particularly in intensive care unit patients by increasing risk of death by 3–7 times ^3, 4^. Specifically, severe AKI requiring renal replacement therapy carries a staggeringly high mortality rate of 50–80% ^3, 5, 6^. Regardless of the underlying cause, no pharmacologic therapies currently exist to prevent or treat AKI. Clinical management remains limited to hemodynamic stabilization, aimed at supporting the kidney’s intrinsic repair processes^5^. There have been many clinical trials of potential pharmacological agents to treat AKI, however none have proven highly effective and safe for patient populations ^7^. Suboptimal pharmacokinetics, off-target toxicity, low efficacy, and molecular instability remain major challenges that prevent the development of effective therapies for kidney disease.

Nanotechnology offers a promising solution and presents new treatment strategies for kidney disorders ^8^. Nanoparticles can be customized as diagnostic or therapeutic delivery vehicles, enhancing drug pharmacokinetics and pharmacodynamics to reduce toxicity, increase efficacy, and improve stability ^9^. They can solubilize poorly water-soluble drugs, enable controlled release, and protect easily degradable biologic cargo. Nanomedicine allows for targeted drug delivery to specific organs and tissues, increasing drug concentration at disease sites and reducing adverse effects on healthy tissues ^10–13^. As a result, nanoparticles have been employed across diverse medical applications, including cancer therapy, iron replacement, fungal or bacterial infections, and genetic liver disorders ^11, 14–16^. Notably, nanoparticles play a crucial role in cutting-edge therapies involving biologics and gene therapy, such as mRNA vaccines. Although nanoparticles hold significant promise, their applications in kidney-related disorders have lagged behind their use in cancer and infectious diseases. Building momentum, emerging therapeutics and recent advances in kidney-targeted nanotechnology are beginning to transform the field. Most kidney-targeted nanoparticles developed in recent years target the kidney passively, based on size and physicochemical properties ^17, 18^. Nanotherapeutics that are actively targeted to the kidney with specific ligands, are mostly targeted to specific markers on epithelial cells ^8, 19^.

Being the gateway to organs, endothelial cells are an attractive cellular target for nanotherapy-based approaches in kidney disease. In the context of AKI, we and others have shown that endothelial cells become pro-inflammatory ^20, 21^, inducing the expression of adhesion molecules, selectins, and chemokines, which together with receptor patterns on immune cells, define the immune profile of inflamed tissue ^22^. Nevertheless, there are significant drug delivery challenges for the translation of these findings into the clinical domain. Previous efforts to facilitate specific targeting of the endothelium have predominantly employed antibody-mediated targeting^23^. While this approach is promising, it requires additional conjugation steps and sourcing of biologics, resulting in an exceedingly complex, costly, and impractical option for the development of drug delivery vehicles. There is therefore a clear need for precise, scalable, and clinically viable strategies to deliver therapeutics specifically to inflamed/injured kidney.

To address this gap, we developed surface-engineered nanoparticles designed for active targeting of inflamed renal endothelial cells. We employed poly (ethylene glycol)-b-poly (propylene sulfide) (PEG-b-PPS) polymer-based nanocarriers, which have been extensively characterized for toxicity, biodistribution, and in vivo stability by our group and others ^24–28^. These nanocarriers can transport both hydrophobic and hydrophilic payloads and can be functionalized with targeting ligands ^25, 26, 28^. We used a micelle (MC) morphology for our nanocarriers, as we have previously shown its superior kidney uptake compared to other organs, as well as compared to other nanocarrier morphologies ^29^. To further increase targeting specificity, we have engineered a unique dual peptide display strategy using peptide amphiphiles for facile, modular incorporation into the self-assembled nanoparticles.

## Materials and Methods

### Chemicals

Unless otherwise stated, all chemical reagents were purchased from the Sigma-Aldrich Chemical Company.

### Synthesis of PEG-b-PPS copolymers

Nanocarriers were fabricated via self-assembly of PEG-b-PPS copolymers. The specific molecular weight ratio of the hydrophilic PEG block to the hydrophobic PPS block was chosen for controlled assembly of a spherical MC morphology. PEG-b-PPS block copolymers were synthesized as previously described ^29^, using PEG thioacetate deprotection by sodium methoxide to initiate anionic ring opening living polymerization of PPS. The reaction was allowed to progress to completion and the PPS was end-capped with benzyl bromide (Supplementary Table 1). The resulting block copolymers (PEG_45_-*b*-PPS_18_) underwent purification by double precipitation in cold diethyl-ether, and their structure was characterized by ^1^H-NMR (CDCl_3_) and gel permeation chromatography ThermoFisher Scientific) using Waters Styragel columns with refractive index and UV-vis detectors in a tetrahydrofuran mobile phase.

### Nanocarrier assembly

MC were self-assembled from the PEG-b-PPS using the thin film hydration method in phosphate-buffered saline (PBS) following previously established procedures ^26, 29^. Briefly, PEG-b-PPS copolymer and fluorescent dye (DiO or DiI, Invitrogen), were dissolved in dichloromethane within 1.8 mL clear glass vials (ThermoFisher Scientific) and subjected to vacuum conditions to eliminate the solvent. The resulting thin films were hydrated in PBS under shaking at 1500 rpm overnight. For in vitro imaging studies, MC suspensions were fabricated using DiO hydrophobic dye, while for in vivo imaging studies, DiI hydrophobic dye was utilized.

### Size and charge of MC formulations

The average nanoparticle diameter (z-size), as well as size distribution, and polydispersity index (PDI) of the MC formulations were assessed by Dynamic Light Scattering (DLS), using a Zetasizer Nano (Malvern Instruments). This analysis was conducted at a concentration of 1 mg/mL in PBS, employing a 4mW He-Ne 633 nm laser. PDI was determined through a two-parameter fit to the DLS correlation data. Zeta potential was determined by performing Electrophoretic Light Scattering (ELS).

### Cryo-transmission electron microscopy (cryo-TEM)

200 mesh Cu grids with a lacey carbon membrane (EMS Cat# LC200-CU-100) were glow discharged using a Pelco easiGlow (Ted Pella) at 15 mA for 30 s under 0.24 mbar pressure, creating a negative charge on the carbon membrane to ensure even liquid sample distribution. 4 µL of sample (5 mg/mL) was applied to the glow discharged grid, blotted for 5 s with a blot offset of +1, and frozen by plunging into liquid ethane using FEI Vitrobot Mark IV. Grids were stored under liquid nitrogen. Grids were then loaded into a Gatan 626.6 cryo transfer holder, images were acquired at –175 °C in a JEOL JEM1400 LaB6 emission TEM at 120 kV, using a Gatan OneView 4k camera. Morphology and size distribution of acquired images was measured using ImageJ.

### Small-Angle X-ray Scattering (SAXS)

Analysis via SAXS was executed at Argonne National Laboratory’s Advanced Photon Source, on the 5-ID beamline, facilitated by the DuPont-Northwestern-Dow Collaborative Access Team (DND-CAT). The technique utilized collimated X-ray beams, having a wavelength of λ = 1.24 Å and an energy of 9 keV. The samples, uniformly prepared to a concentration of 5 mg/mL, were examined using a flow cell system situated in-vacuum, enclosed by quartz capillaries with a 1.6 mm wall thickness. We obtained scattering data over a momentum transfer range (q-range) from 0.0015 to 0.08 Å⁻¹. The setup involved positioning the sample around 8.5 meters from the detector, with each exposure lasting 5 seconds. Calibration of the instrument was conducted using silver behenate and a gold-coated silicon grating with a 7200 lines/mm density. The momentum transfer vector q is calculated as q = (4π/λ), where 2θ is the angle of scattering. Data processing, encompassing reduction and buffer subtraction, was handled with the BioXTAS RAW program. Furthermore, the SasView 5.0.5 software facilitated the fitting of models to the scattering data ^30^.

### Synthesis of targeting peptide constructs and incorporation into MC

Constructs comprised of peptide/PEG/Palmitoleic acid were synthesized by the Simpson Querrey Institute (SQI) peptide synthesis core. Fmoc-N-amido-dPEG_24_-amido-dPEG_24_-acid (Quanta Biodesign) was purchased for the synthesis of these peptide constructs. Conventional Fmoc solid-phase peptide synthesis was employed for the generation of PG_24_ × 2 (PG_48_) peptides, on a 0.1 mmol scale of each targeting sequence: inflamed endothelium peptide (CYNTTTHRC) and kidney peptide (CLPVASC). Information regarding each synthesized peptide-construct is presented in Supplementary Table 2. The peptide was synthesized with an amide on the c-terminus, and underwent initial purification in its linear form, prior to disulfide cyclization and re-purification. The purified molecules were prepared in acetonitrile/water with 0.1% trifluoroacetic acid.

Targeting peptide constructs were dissolved in dimethyl sulfoxide and added to MC aliquots at the desired molar ratio (1%, 2.5% or 5% peptide/polymer) allowing precise control over the density of peptide modifications on the MC. Non targeted MC controls (lacking peptides) were included in all uptake studies. The various formulation vials were left to roll overnight. All formulations were prepared under sterile conditions and underwent purification using a Sephadex LH-20 gravity column with a PBS mobile phase. The peptide incorporation into purified MC, was confirmed by MALDI-TOF-MS performed after MC purification, depicting specific dominant peaks that corelate with the extracted mass spectra for the two peptide constructs when these were synthesized.

### Cell Culture

Human primary pulmonary artery endothelial cells (HPAEC) were purchased from ATCC and cultured in endothelial cell growth medium-2 (EGM-2, Lonza), supplemented with the EGM-2 SingleQuot Kit (Lonza). Cultivation was carried out in cell culture flasks and multi-well plates pre-coated with 0.1% gelatine. The cell culture environment was maintained at 37°C with 5% CO_2_, and the cell media was routinely refreshed. For cell passage, a standard trypsinization procedure was employed, and cells were split when reaching 75-80% confluency. HPAEC used in all experiments were limited to passage ≤7. The 3-(4,5-dimethylthiazol-2-yl)-2,5-diphenyltetrazolium bromide (MTT) assay was used to assess the viability of HPPAEC following treatment with MCs as previously described ^31^.

### In vitro cell uptake studies

Hypoxia-reoxygenation studies were used to mimic an ischemia-reperfusion injury (IRI) induced inflammatory state. HPAEC were seeded at a density of 20,000 cells/well in 96-well plates and allowed to adhere overnight at 37 °C, with 5% CO2. Subsequently, cells were incubated for 14.5-19 h in a hypoxia chamber (Coy Laboratory Products) at 0.5% O_2_, 37°C in a 5% CO_2_ humidified environment. During reoxygenation, cells were treated with 1 ng/ml IL1β for 7-11 h. Following this, cells were treated with specified DiO-loaded MC formulations (5 mg/mL polymer), differing by the molar ratio of the targeting peptide [0%, 1%, 2.5% and 5% of the inflamed endothelial cell (IEC) targeting peptide], with the additional incorporation of the kidney targeting peptide (1% or 5%). Cells were incubated for 2 h at 37°C, 5% CO_2_ and thereafter were washed three times with PBS to eliminate free MC. Each experimental set included untreated cells and a PBS-treated group, with three biological replicates per treatment group (n = 3). MC uptake was quantified using a fluorescent microscope (Leica DM IL LED). The median fluorescence intensity (MFI) above the PBS-treated background was calculated to subtract cellular autofluorescence contributions to the measured values.

### Mice

C57BL/6J mice were purchased from the Jackson Laboratory (BarHarbor, ME), and subsequently bred and housed in the Center for Comparative Medicine at Northwestern University. The mice were kept in clear cages in pathogen-free housing rooms at 21°C with a 12-h light/12-h dark cycle and were provided with a standard diet. All animal experimental procedures adhered to protocols approved by the Northwestern University Institutional Animal Care and Use Committee (IACUC). In each experiment, mice were randomly allocated to experimental groups.

### In vivo uptake studies

We employed a previously established and well-characterized unilateral renal IRI model ^20^ to evaluate the targeting effects of our formulations in vivo. Female mice aged 17.5 to 22.5 weeks were utilized for these experiments. For the IRI model, mice were anesthetized with an intraperitoneal injection of ketamine (90–120 mg/kg) and xylazine (10 mg/kg). A small midline abdominal incision was made, and the left renal pedicle was occluded with a microaneurysm clamp, while the right kidney served as an internal control. The abdominal incision was temporarily sutured partially closed, and body temperature was monitored by rectal probe and maintained at 37°C using a heating pad. After 30 minutes, the clamp was removed, and reperfusion was visually confirmed. The abdominal facia was closed with a 6-0 suture, and Michel miniature clips were used to close the skin. Mice were kept on a heating pad until recovery from the anesthesia and then returned to the animal housing rack. 26-31 h post-surgery, mice were randomly assigned to three groups (4 mice/group). All mice were intravenously injected (retro-orbitally) with 75 microliters of formulation. The two treatment groups received non-targeted MCs (with no peptide) vs targeted MCs (with a 1% molar ratio of each targeting peptide). The MC concentration was 17 mg/ml, and they were loaded with 0.3% weight: weight DiI. The control group received a retro-orbital injection of PBS (75 microliter) at the same time point after IRI.

18-20 h post-injection, mice were sacrificed by CO_2,_ and organs of interest (lungs, liver, spleen, heart and kidneys) were harvested in PBS filled Petri dishes. In-Vivo Imaging System (IVIS) Lumina scans were performed (Center for Advanced Molecular Imaging, Northwestern University) with λexc = 745 nm, λem = 810 nm, exposure time = 2 s and f/stop = 2. Organ imaging was performed ex-vivo, simultaneously comparing mouse organs from all three groups in each IVIS scan, to best assess relative radiance efficiency. The experiment was performed on two separate occasions (total of n=8 per group).

### Statistical analysis

Statistical analyses were performed using Prism software (version 10.1.2; GraphPad Prism Software, LLC). Two-group comparison was performed by unpaired 2-tailed Student’s *t* test with Welch’s correction. Multigroup comparison was performed by 1-way ANOVA with Turkey’s multiple-comparison test

## Results

### Synthesis and characterization of the peptide targeted nanocarriers

Because our prior studies demonstrated that PEG-b-PPS micelles (MC) accumulate in the kidney ^29^, we chose them for our active targeting studies. First, PEG_45_-*b*-PPS_18_ block copolymers were synthesized and self-assembled into spherical MC following established procedures (**Figure 1a**). To enhance organ- and injury-specific delivery, we aimed to engineer MC displaying two targeting peptides: one with kidney tropism and another targeting inflamed vasculature (**Figure 1a&b**). To achieve this, we incorporated peptides previously reported to have preferential distribution to kidney and targeting vascular inflammation, respectively. Kidney specific targeting has been approached with several peptides, mostly using epithelial targets ^8^. As the CLPVASC (**Figure 1b**) sequence has been found to be specific to kidney tissue without being specific to epithelial cells ^32^ and has successfully improved kidney targeting of nanocarriers by others ^33^, we employed it in our targeting system to enhance kidney delivery. Several peptide sequences binding specifically to inflamed endothelium were previously identified using phage display^34^. From these described sequences, we selected CYNTTTHRC (**Figure 1b**) as a targeting moiety due to its maximal specificity to inflamed endothelial cells (IEC) in kidneys vs other organs, such as liver, heart, and lungs. The peptide (hereinafter referred to as “IEC”), containing NTTTH domain, was found to be specific to inflamed endothelium, while showing no affinity to endothelium without concomitant inflammation. The peptide is homologous to cell surface proteins Scube 1 and 2, which have previously been functionally associated with endothelial inflammation ^35^. Peptide constructs were designed, displaying each of these two peptides (**Figure 1b**). We have previously shown that the use of PEG_48_ spacers to mount targeting peptides improved MC targeting compared to shorter PEG spacers ^24^. Due to this, the peptide constructs in our study were synthesized with 48-unit PEG spacers (slightly longer than the PEG component of the block copolymer). The two targeting peptides were synthesized using standard Fmoc solid-phase peptide synthesis, and the resulting products were of high purity (≥95% purity). Targeting peptides were incorporated using lipid-anchored constructs, a method previously shown to preserve micelle size and structural integrity ^26^. Peptide lipid constructs were embedded into PEG-*b*-PPS MC nanocarriers at a 1%, 2.5% or 5% molar ratio (peptide/polymer) and were purified through a lipophilic Sephadex column to remove any unembedded peptide. The resulting MC formulations were monodisperse (PDI<0.1) with an average diameter of ∼23.5−23.8 nm (**Table 1**). ELS analysis demonstrates a zeta potential indicative of a neutrally charged surface (zeta potential −6.2 to −2) for all nanocarriers in the presence and absence of peptide (**Table 1**). Dominant peaks of 3183, and 3588 DA were visible in the extracted mass spectra for CLPVASC, and CYNTTTHRC, respectively, and these mass differences were consistent with the differences in the mass of the construct-specific peptides (**Figure 1c**). The incorporation of peptide did not disrupt the spherical morphology expected for PEG-*b*-PPS MCs, as demonstrated by morphological analysis using cryogenic transmission electron microscopy (cryo-TEM; **Figure 1d**; see the “methods” information for cryo-TEM procedures). The structural integrity of MC upon peptide construct incorporation was further investigated using DLS (**Table 1**) and SAXS (**Figure 1e and Table 2**). SAXS analysis revealed that the peptide conjugation did not alter the micellar structure, as indicated by the fitting of the SAXS data to a monodisperse population model with an excellent fit (χ² << 1), suggesting that the size distribution is narrow, and the MCs are well-formed. Complementary DLS measurements showed a hydrodynamic radius that was approximately 5 nm larger than that determined by SAXS (**Table 1** and **Table 2**). This is consistent with expectations, as DLS measures the hydrodynamic size, which includes the MC along with its solvation layer (PEG corona), while SAXS measures the core size of the MC itself ^36^. There were not significant differences in MC size between blank and peptide-displaying formulations by DLS or by SAXS (**Table 1** and **Table 2**). These results are consistent with our previous findings that incorporation of targeting peptide via lipid anchoring does not alter the size or structure of MCs ^26^. Furthermore, they confirm that the addition of peptide constructs at a maximal concentration of 5% molar ratio for each peptide (10% total) does not disrupt their structural integrity, while MC remain monodisperse with a consistent size distribution after modification. Finally, peptide incorporation into MC was verified by MALDI TOF-MS (**Figure 1f**).

**Figure 1.**
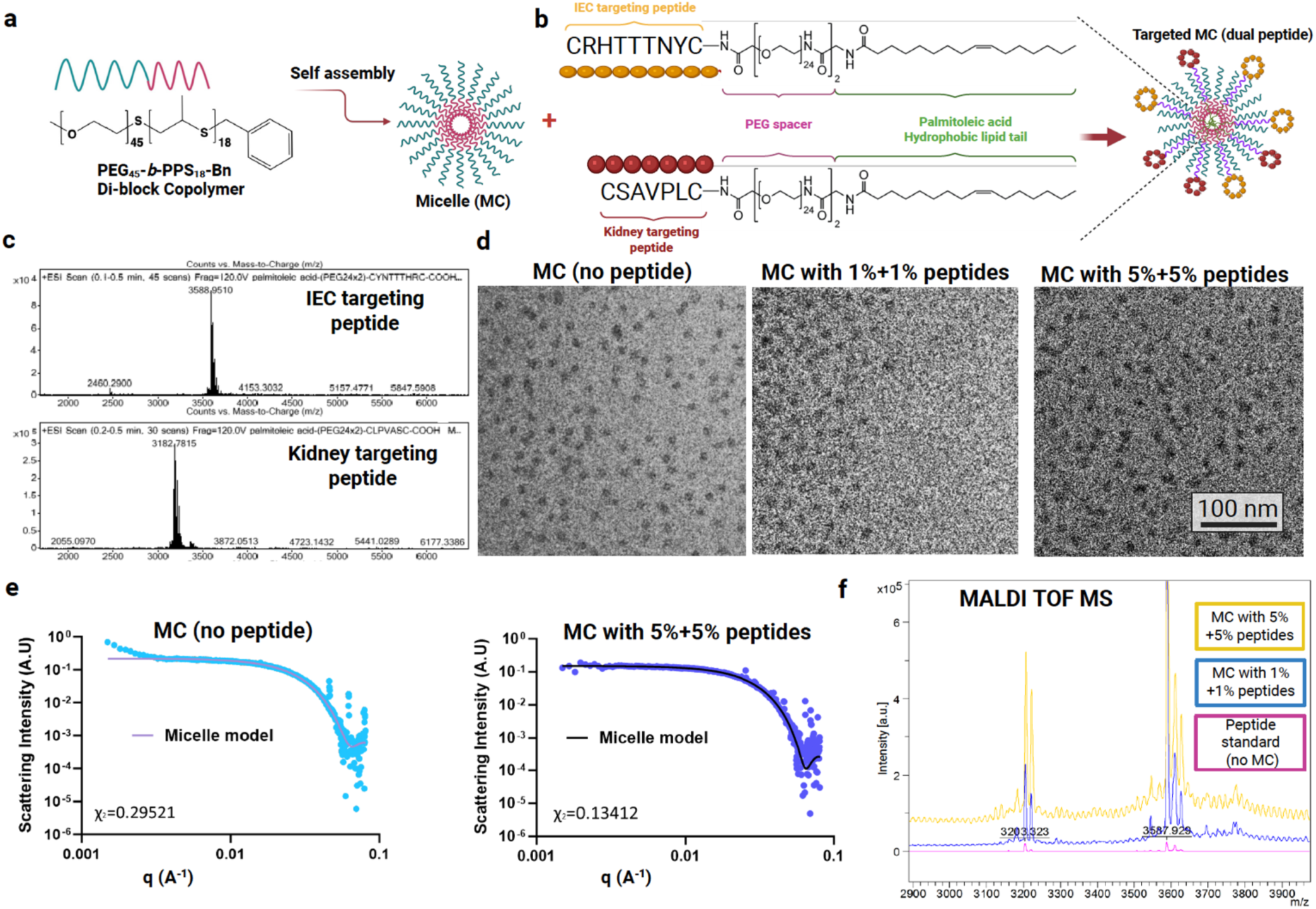
Schematic and characterization of targeted nanoparticles. (a) Schematic representation and chemical structure of poly(ethylene glycol)-block-poly(propylene sulfide) (PEG-b-PPS) copolymers which were self-assembled into micelles (MC). (b) MC are combined with the designed dual targeting peptide (IEC + Kidney) constructs anchored to the MC by lipid tails. (c) Lipid anchored peptide constructs were synthesized of the form [palmitoleic acid]-[PEG_48_ spacer]-[targeting peptide] (listed C-terminus to N-terminus). The constructs differed by the targeting peptide; Top: Inflamed endothelial targeting peptide; Bottom: Kidney targeting peptide. The targeting peptides are both cyclic via disulfide bonds. (d,e) MC morphology was verified by cryogenic electron microscopy (Cryo-TEM) (d) and by Synchrotron small angle x-ray scattering (SAXS) (e). The structure remained consistent both when non-targeted (i.e., without peptide) and targeted (i.e., with the peptide constructs). The magnification is 10,000×, and the scale bar is 100 nm for cryo-TEM micrographs. In all cases, SAXS was performed using synchrotron radiation and a core−shell model (solid line) was fitted to the data (blue dots). χ2 ≪ 1.0 was obtained for all model fits (a good fit is indicated by χ2 < 1.0). (f) MALDI-TOF was used to confirm that peptide constructs were embedded in MCs, with the mass indicated as 3201.

**Table 1.** Physicochemical Characteristics of PEG-b-PPS Micelles Displaying Targeting Peptide Constructs.

| Formulation | DLS<sup>a</sup> |  | ELS |
| --- | --- | --- | --- |
|  | *D* [nm] | PDI | zeta potential [mV]<sup>b</sup> |
| Micelle (no peptide) | 23.47 | 0.144 | -6.2 ± 6.2 |
| Micelle with 1%+1% targeting peptides | 23.81 | 0.181 | -2.0 ± 1.0 |
| Micelle with 5%+5% targeting peptides | 23.68 | 0.198 | -3.2 ± 0.6 |

<sup>a</sup> Z-average hydrodynamic diameter (*D*) and PDI determined by DLS (n = 3)
<sup>b</sup> Mean zeta potential ± standard deviation measured by ELS (n = 3)

**Table 2.** Physical Characteristics of PEG-b-PPS Micelles Displaying Targeting Peptide Constructs by SAXS^a^.

| Formulation | *D*<sub>Total</sub> [nm] | *R*<sub>Core</sub> [nm] | $\chi^2$ |
| --- | --- | --- | --- |
| Micelle (no peptide) | 18.1 | 6.9 | 0.295 |
| Micelle with both targeting peptides | 17.9 | 6.9 | 0.134 |

<sup>a</sup>SAXS conducted using synchrotron radiation. The fitted parameters include the total diameter (*D*<sub>Total</sub>) and core radius (*R*<sub>Core</sub>), along with the chi-squared ( $\chi^2$ ) value indicating the final model fit. Further details can be found in the Materials and Methods section.

### Peptide targeting increases nanocarrier uptake by inflamed HPPAECs in vitro

We first tested whether the synthesized PEG-b-PPS micelles had any cytotoxicity to Human Primary Pulmonary Artery Endothelial Cells (HPAEC). HPPAEC were incubated with MC loaded with fluorescent die, at concentrations up to 18.9 mg/dl. Growth reduction compared to PBS was less than 15% for even the highest concentration (data no shown), which is consistent with our previous findings. The targeting efficacy of the targeted MC was examined in vitro using HPAEC stimulated by Hypoxia/ Reoxygenation with IL1β, hereinafter referred to as “inflamed HPAEC”. MC loaded with DiO hydrophobic fluorescent dye were incubated for 2 h with inflamed HPPAEC and their uptake was evaluated using a fluorescent microscope (**Figure 2a**). First, we assessed the effect of the IEC targeting peptide by comparing uptake of MC with no targeting peptide to MC with the peptide construct. Fluorescence analysis demonstrated that the IEC peptide incorporation into MC directly increased MC uptake by HPPAECs, compared to non-targeted MC (containing no peptide) (**Figure 2b** and **Supplementary Figure 1**). Aiming to achieve organ-specific targeting, we combined a Kidney-targeting peptide at 1% or 5% density in MC with 1% IEC-peptide density and examined whether this addition interferes with the IEC-targeting effect described above. Our in vitro studies showed that the uptake of MC by inflamed endothelial cells was better preserved in the presence of 1% kidney peptide as opposed to 5% kidney peptide (**Figure 2c**).

**Figure 2.**
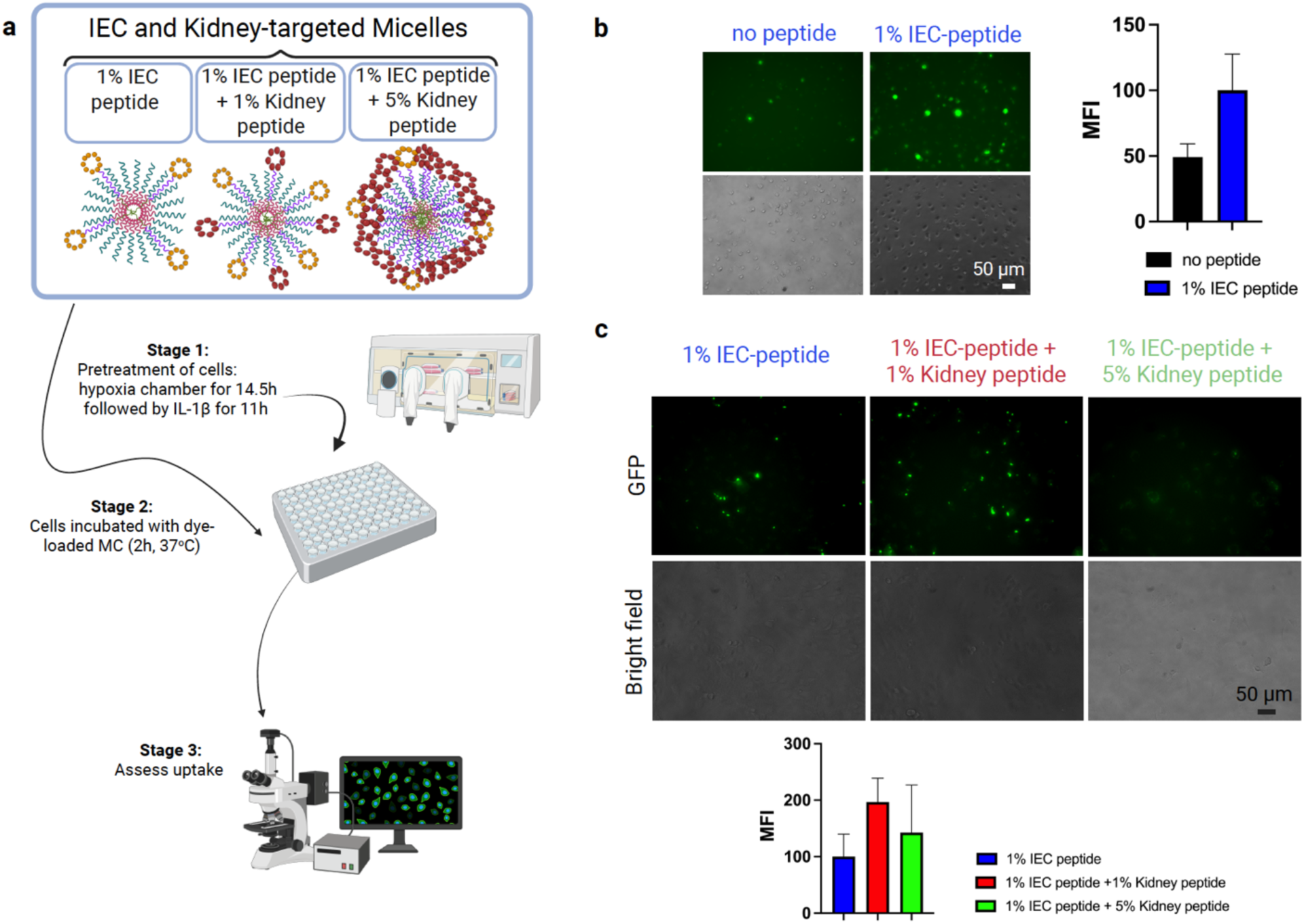
Targeted nanoparticle uptake by inflamed human endothelial cells in vitro. (a) Schematic of PEG-b-PPS MC formulations and the cellular uptake study. Following hypoxia-reoxygenation and exposure to IL-1β, inflamed HPPAECs were incubated at 37° C for 2 h with DiO loaded MCs that were incorporated with various amounts of targeting constructs. The constructs varied by molar ratio of peptide to MC polymer (1% of inflamed endothelial cell (IEC) peptide and addition of 1% or 5% kidney peptide). After 2 h incubation, cells were washed 3x and imaged by a Leica fluorescent microscope. (b &c) Shown are representative images of the MC uptake across the indicated MC formulations. Median fluorescence intensity (MFI) was measured to quantify uptake. The use of dual peptide targeting with the addition of a Kidney specific peptide maintains the increased uptake, without reducing the IEC peptide’s targeting effect.

### Dual peptide targeting increases MC distribution to the inflamed kidney

To examine whether out dual peptide approach enhances delivery to the inflamed kidney, we used a mouse model of unilateral IRI. In this model, one kidney is subjected to IRI, induced by renal pedicle clamping for 30 minutes. The contralateral kidney remains intact and serves as an internal control. We and others have extensively characterized this model and have shown that IRI induces inflammation in various kidney cell types, including endothelial cells ^31, 37, 38^. Notably, even remote organs, such as the lungs, heart, liver and spleen, develop inflammatory responses secondary to unilateral kidney IRI ^39–41^. Because MC with peptide density of 1% for the IEC- and kidney-targeting peptide showed targeting capacity in vitro, we employed them as the targeted nanocarriers of choice in our animal studies. Specifically, mice underwent unilateral renal ischemia-reperfusion injury (IRI) and received intravenous injections (retro-orbital) of either targeted MC, non-targeted MC, or PBS at 25-31 hours post-IRI (corresponding to Day 1 post IRI, n = 8 per group). Targeted MC displayed both IEC- and Kidney-targeting peptides (1% molar ratio each) and were loaded with DiI (0.3% w/w; 17 mg/mL). Mice were sacrificed at Day 2 post-IRI (18–20 hours after injection), and major organs (kidneys, liver, lungs, heart, spleen) were harvested (**Figure 3a**). Ex vivo biodistribution was assessed using IVIS imaging, comparing relative radiant efficiency across treatment groups (**Figure 3b**).

**Figure 3.**
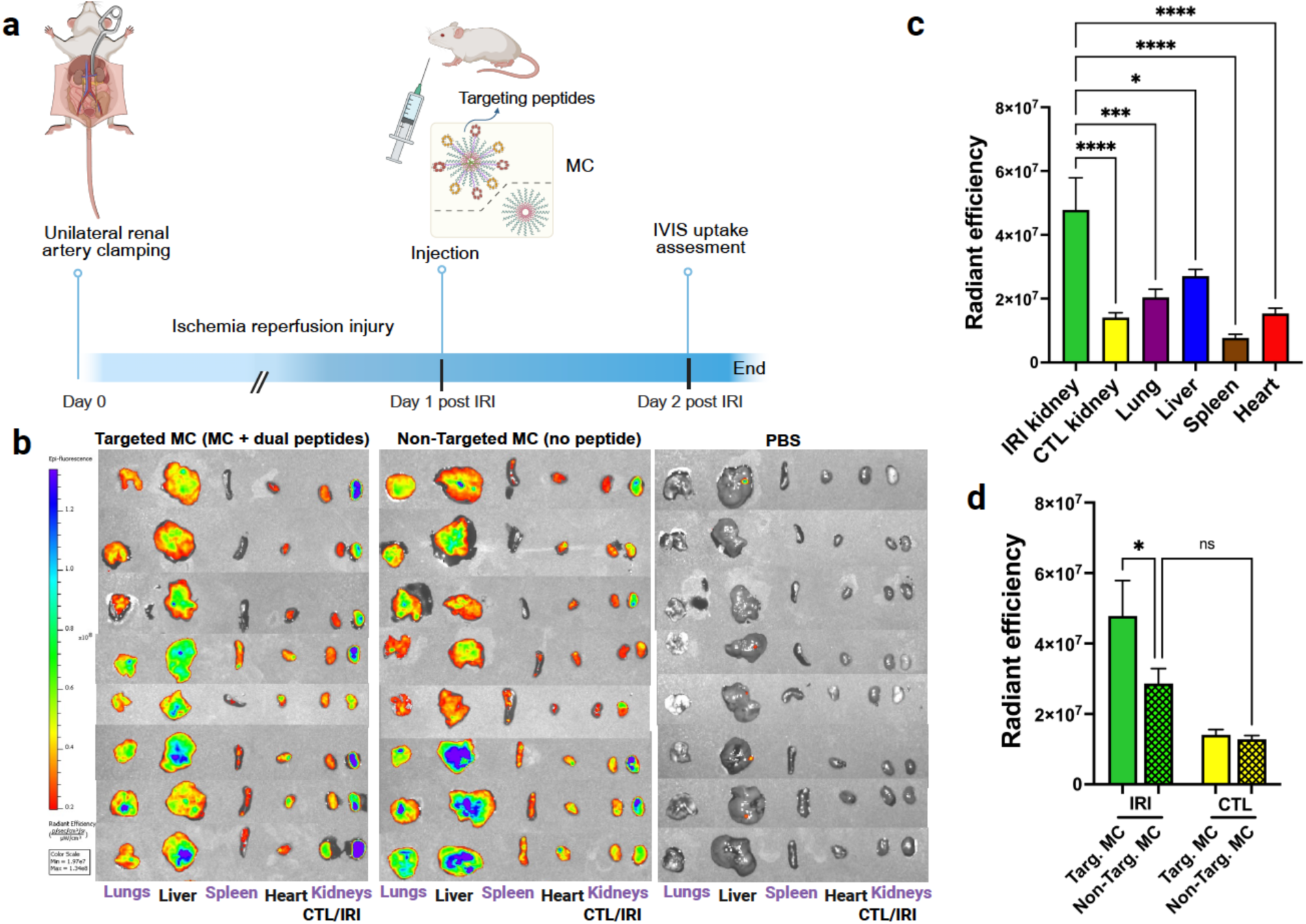
Nanocarriers presenting dual targeting peptides selectively enhance uptake by inflamed kidneys and concurrently reduce off-target uptake in a murine model of unilateral renal ischemia reperfusion injury. (a) Illustrative experimental overview: The targeting of PEG-b-PPS MCs displaying dual targeting peptides compared to non-targeted MCs (with no targeting peptide) was evaluated in a model of IRI induced by unilateral renal artery clamping. Targeted and non-targeted nanocarriers were injected systemically via mice retrobulbar plexus (n=8) for each group, at 26 to 30.5 h after renal artery clamping, and compared to a third control group, injected with PBS. Nanocarrier uptake was assessed ∼19 h after injection using ex vivo organ IVIS. (b) Organ uptake represented by IVIS. (c-d) Radiant efficiency was calculated from IVIS software, and each organ uptake was calculated. In the context of IRI, targeted MC showed significantly increased uptake by the injured/inflamed kidney compared to the contralateral kidney and other peripheral organs tested; lung, liver, spleen and heart (c). Targeted MC showed significantly increased uptake in the IRI kidney compared to non-targeted MC (d). For (c-d), Data shown are mean ± SEM, statistical significance was determined by 1-way ANOVA with Turkey’s multiple-comparison test. CTL; contralateral kidney, IRI; kidney subjected to ischemia-reperfusion injury, Targ; Targeted, Non-Targ: Non-targeted. *, p < 0.05; **, p<0.01, ***, p<0.001; ****, p<0.0001; ns: non-significant.

Targeted MC exhibited significantly increased uptake by 3.4-fold in the IRI kidney compared to the contralateral control kidney (n=8, p<0.0001) (**Figure 3c**). For the non-targeted MC, injury increased MC uptake in the IRI kidney compared to contralateral without reaching statistical significance though (**Figure 3d**). Furthermore, the IRI kidneys displayed significantly augmented uptake of targeted MC compared to non-targeted MC by 67% (**Figure 3d**), whereas contralateral kidneys showed no difference in this context. Upon evaluation of distribution to off-target organs, dual targeting significantly reduced MC delivery within the other tested organs compared to the IRI kidney (**Figure 3c**). For instance, compared to the IRI kidney, the uptake of targeted MC was lower in the lungs by 2.3-fold (P<0.0001). Taken together, these findings demonstrate that dual peptide targeting of MC can significantly enhance distribution to the inflamed kidney in a mouse model of unilateral kidney IRI.

## Discussion

Over the past decade, various strategies have been explored to improve nanoparticle delivery to the kidney, most relying on passive mechanisms such as particle size, charge, and surface chemistry. While these factors can influence renal accumulation, most systems lack consistent kidney specificity in vivo. Active targeting using surface ligands, most commonly peptides or antibodies, offers greater specificity. Peptides, in particular, are attractive due to their small size, ease of synthesis, biocompatibility, and tunable chemistry, and recent advances in phage display and computational modeling have expanded the repertoire of peptides with kidney and cell-type specificity ^42^. Compared to antibodies, peptides are more amenable to scalable manufacturing and modular nanocarrier integration, although challenges such as in vivo stability and moderate binding affinity remain areas for continued optimization ^43^. In our study, we used PEG-b-PPS micelles, previously shown to exhibit favorable biodistribution ^25, 28^ and immunological inertness ^27, 29^, as a base for dual-functionalized nanocarriers that co-display kidney- and endothelial inflammation-specific peptides. This approach addresses current limitations in spatial and pathophysiological precision and offers a modular strategy to improve therapeutic localization in AKI.

In our previous work, by performing comprehensive ex-vivo analysis of fluorescent nanocarriers within various mouse organs 24 h after systemic injection, we found that PEG-b-PPS MC preferably accumulate in the liver and kidneys ^29^. Here, we designed MC to incorporate two different specific targeting peptides simultaneously on the same particle. These peptides were previously reported and identified through traditional high-throughput screening methods. Our molecular characterization demonstrated that this unique targeting technique did not alter the size, charge, or structure of the polymeric MC from their well-studied basic form. In vitro studies confirmed that optimal uptake by inflamed endothelial cells was not hindered by the addition of the kidney-specific peptide. Looking ahead, advances in AI-based tools such as AlphaFold present exciting opportunities to streamline and expand the discovery of high-affinity targeting peptides through virtual screening ^44^.

Our in vivo studies in a murine model of kidney IRI demonstrate that surface-engineered PEG-b-PPS micelles functionalized with dual-specific peptides enhance delivery to the injured kidney. This dual-targeting strategy, which combines organ-level and injury-state specificity, offers a modular and effective approach for improving the spatial precision of therapeutic delivery in AKI. While biodistribution analyses for the dual peptide-decorated MC revealed significantly greater accumulation in the injured kidney compared to other organs, further optimization will be important to improve relative targeting efficiency and minimize unintended delivery. Techniques such as indirect targeting could be combined with the nanocarriers developed here to minimize their off-target accumulation via non-specific mechanisms of pinocytosis ^45^. Nonetheless, the ability to direct nanocarriers to the site of injury avoiding systemic undesired distribution underscores the translational potential of this platform. Given the clinical limitations of current systemic therapies for kidney diseases, including widespread toxicity and inadequate renal targeting, our findings provide proof-of-principle for a peptide-based delivery system that could be adapted for diverse therapeutic payloads. As nanomedicine advances, such rationally designed, and biologically informed platforms may help bridge the gap between precision delivery and clinical application in kidney therapeutics.

In summary, the dual peptide-based delivery approach established here offer a proof-of-principle for precise targeting of inflamed kidney in AKI. This strategy holds strong potential to improve therapeutic outcomes in diverse clinical settings and may help catalyze further translational advances in kidney nanomedicine.

## Conclusions

Our work has shown that a dual peptide-based targeting strategy can enhance the delivery of nanocarriers to the post-ischemic kidney. This represents a significant technological advance enabling improved targeted delivery of novel kidney therapeutics.

## Author contributions

B.Y.B. and R.T. designed the research with input from E. A. S., S.E.Q and P.P.K. B.Y.B, S.H.S, R.T. and S.A. performed experiments with the assistance of S.A.Y. B.Y.B., E. A. S. and P.P.K wrote and revised the manuscript. All authors have given approval to the final version of the manuscript.

## Data availability

The authors confirm that the data supporting the findings of this work is available within the article. The raw data is also accessible from the corresponding author upon reasonable request.

## Acknowledgements

The reported study was supported by a P&F grant from the Northwestern University George M. O’Brien Kidney Research Core Center (NUGoKidney) (P30 DK114857) and the National Institute of Health grants U54DK137516 (SEQ, PPK), R01DK115850 and R01DK1326721. This work benefited from the use of the SasView application, originally developed under NSF award DMR-0520547. SasView contains code developed with funding from the European Union’s Horizon 2020 research and innovation program under the SINE2020 project, grant agreement No 654000. BB was a recipient of a post-doctoral fellowship from the NU KUH training grant (TL1DK132769). The content is solely the responsibility of the authors and does not necessarily represent the official views of the National Institutes of Health.

## Supplementary Data

**Table S1.**
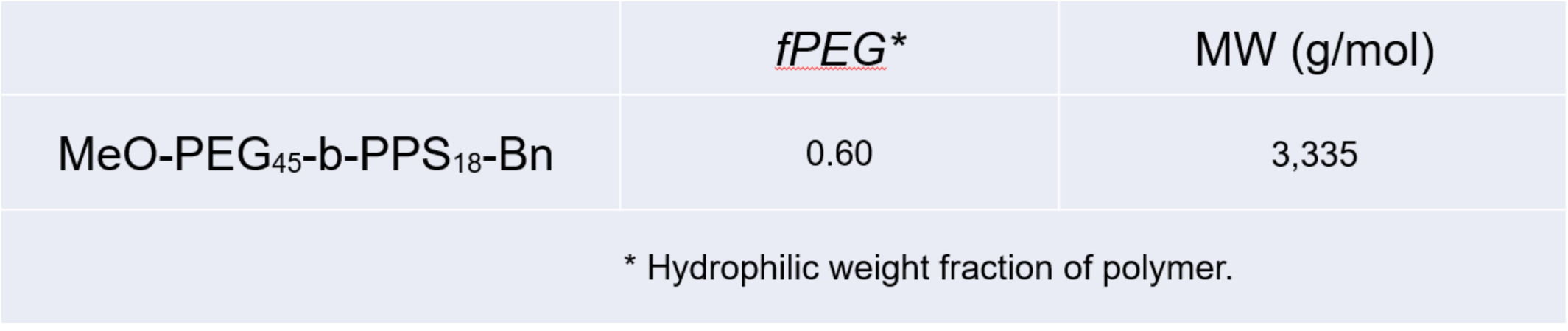
PEG-*b*-PPS Micelle (MC) diblock copolymer.

**Table S2.** Targeting peptide constructs.

| Construct* | Lipid Anchor** | PEG Spacer*** |  | Peptide primary structure‡ | MW (g/mol) | Purity‡‡ |
| --- | --- | --- | --- | --- | --- | --- |
|  |  | Units | Length (Å) |  |  |  |
| PG<sub>48</sub> | PA | 48 | 174.058 | CLPVASC | 3182 | >95% |
| PG<sub>48</sub> | PA | 48 | 174.058 | CYNTTTHRC | 3588 | >95% |

\* All peptide constructs are of the form [Lipid]-[PEG<sub>x</sub> spacer]-[Cyclic peptide].
\*\* Palmitoleic acid (PA).
\*\*\* The spacer length is presented as the number of PEG units.
The peptide construct with a 48-unit PEG spacer was synthesized using Fmoc-PEG24x2.
‡ Peptide primary structure is listed N-terminus to C-terminus.
‡‡ Peptide purity determined using LC-MS.

---

**Figure S1.**
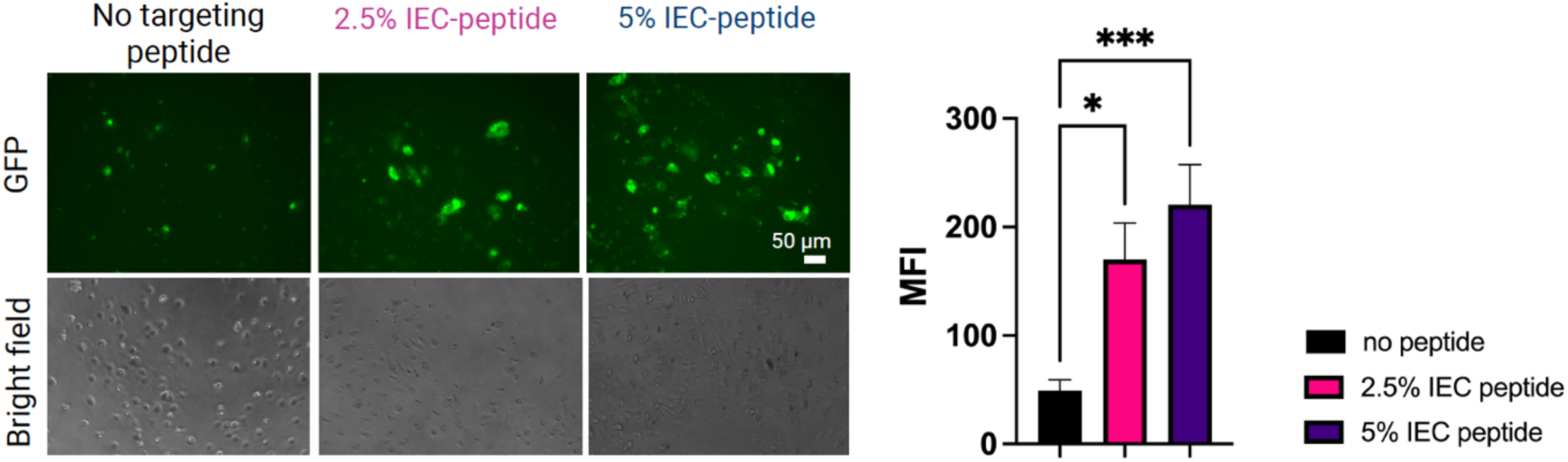
MC functionalized with IEC-peptide demonstrate increased uptake by inflamed human endothelial cells. Shown are representative images of the MC uptake across the indicated MC formulations. Median fluorescence intensity (MFI) was measured to quantify uptake. Statistics were determined using one-way ANOVA with Sidak correction for multiple comparisons. *, P<0.05; ***, P<0.001.

